# Identifying the Brain Circuits that Regulate Pain-Induced Sleep Disturbances

**DOI:** 10.1101/2024.12.20.629596

**Authors:** Nicole Lynch, Roberto De Luca, Richard L Spinieli, Enrico Rillosi, Renner C Thomas, Samuel Sailesh, Nishta Gangeddula, Janayna D Lima, Sathyajit Bandaru, Elda Arrigoni, Rami Burstein, Stephen Thankachan, Satvinder Kaur

## Abstract

Pain therapies that alleviate both pain and sleep disturbances may be the most effective for pain relief, as both chronic pain and sleep loss render the opioidergic system, targeted by opioids, less sensitive and effective for analgesia. Therefore, we first studied the link between sleep disturbances and the activation of nociceptors in two acute pain models. Activation of nociceptors in both acute inflammatory (AIP) and opto-pain models led to sleep loss, decreased sleep spindle density, and increased sleep fragmentation that lasted 3 to 6 hours. This relationship is facilitated by the transmission of nociceptive signals through the spino-parabrachial pathways, converging at the wake-active PBel^CGRP^ (parabrachial nucleus expressing Calcitonin Gene-Related Peptide) neurons, known to gate aversive stimuli. However, it has never been tested whether the targeted blocking of this wake pathway can alleviate pain-induced sleep disturbances without increasing sleepiness. Therefore, we next used selective ablations or optogenetic silencing and identified the key role played by the glutamatergic PBel^CGRP^ in pain-induced sleep disturbances. Inactivating the PBel^CGRP^ neurons by genetic deletion or optogenetic silencing prevented these sleep disturbances in both pain models. Furthermore, to understand the wake pathways underlying the pain-induced sleep disturbances, we silenced the PBel^CGRP^ terminals at four key sites in the substantia innominata of the basal forebrain (SI-BF), the central nucleus of Amygdala (CeA), the bed nucleus of stria terminalis (BNST), or the lateral hypothalamus (LH). Silencing of the SI-BF and CeA also significantly reversed pain-induced sleep loss, specifically through the action on the CGRP and NMDA receptors. This was also confirmed by site-specific blockade of these receptors pharmacologically. Our results highlight the significant potential for selectively targeting the wake pathway to effectively treat pain and sleep disturbances, which will minimize risks associated with traditional analgesics.

**One sentence summary:** Parabrachial CGRP neurons regulate awakenings to pain.

## Introduction

Estimates suggest that more than 1 in 5 adults in the United States experience chronic pain, and over 70% of these individuals report ongoing sleep disturbances [1,2]. There is a strong bidirectional relationship between sleep and pain: chronic pain not only reduces sleep quality, but poor sleep can also lead to increased pain and predict higher pain levels [3]. While shorter sleep durations are detrimental, frequent awakenings are the most common complaint among those with chronic pain [4]. A recent study demonstrated that in a chronic pain model, there was significantly increased sleep fragmentation but no reduction in total sleep time [5]. Moreover, sleep deprivation has been shown to heighten pain sensitivity, which can be restored to normal levels with adequate sleep recovery and even improved with wake-promoting agents like caffeine, despite the accumulated sleep debt from deprivation [6,7]. To further examine the interactions between pain and sleep, in this study we investigated the brain circuits that relay pain signals and disrupt normal sleep in two different acute pain models by targeting manipulations of specific cell types and their associated pathways.

The parabrachial nucleus (PB), a wake-active node in the brain stem, receives nociception signals directly from the dorsal horn as part of the spine-ponto-amygdaloid tract [8–11], and mediates arousal in response to numerous aversive sensory stimuli [12]. Activation of glutamatergic neurons in the PB promotes hyperalgesia and neuropathic pain in mice with a chronic peroneal nerve injury, while their inactivation provides analgesia [13]. However, the neuronal population in the PB is heterogeneous and diverse, both functionally and phenotypically, and has distinct projection patterns that are spatially separated [13–16]. In exploring the functionality of the different PB subpopulations, we found that cortical arousal in response to hypercapnia could be prevented by inhibiting a selective cluster of glutamatergic neurons located in the external lateral PB (PBel) that express calcitonin gene-related peptide (PBel^CGRP^)[16]. While another population, that expresses the transcription factor Fork-head box protein (FoxP2) in the centro-lateral part of PB and the Kӧlliker fuse, regulates respiration and respiratory effort during apneas [17]. Subsequently, other studies also identified that PBel^CGRP^ neurons are not only activated by acute painful stimuli [12] but their activity remains amplified for 5 weeks post neuropathic or inflammatory pain exposure, in correlation with increased pain metrics[18].

Collectively, these findings led us to hypothesize that PBel^CGRP^ neurons function as a relay node for transmitting pain stimuli via their forebrain projections to mediate cortical arousals. To investigate this, we studied the effects of pain-induced sleep modifications either by genetic ablation or by the optogenetic inhibition of the PBel^CGRP^ neurons in two acute pain models: inflammatory pain and opto-pain. To identify the arousal pathways associated with the PBel^CGRP^ neurons that may mediate these sleep effects, we also optogenetically inhibited their terminals at the four mains sites that receive moderate to substantial inputs from PBel^CGRP^ neurons[16]: the substantia innominata of the basal forebrain (SI-BF) area, the central nucleus of the amygdala (CeA), the bed nucleus of the stria terminalis (BNST), and the lateral hypothalamus (LH). Some of these sites have been previously indicated as important in pain transmission; however, their roles in modulating pain-related sleep loss have not yet been thoroughly studied [8,15,19]. Finally, we tested local pharmacological blocking using either CGRP or glutamate receptor blockers at two of the main terminal sites (SI- BF and CeA), where optogenetic inhibition had the most significant effects on reversing sleep loss due to acute inflammatory pain (AIP). Our findings provide important insights into the complexity of pain-induced arousal circuitry at a cell-specific level.

## Results

### Acute pain models

We used two models of acute pain stimulus to study the role of the PBel^CGRP^ as relay neurons for pain-induced arousal. Both of these models produced significant and comparative sleep loss and fragmentation.

*1. Acute inflammatory pain (AIP) model:* In this model, animals received a single subcutaneous injection of either 5% formalin or 0.9% buffered sterile saline (25 µl) in one of the hind paws during the early light phase on the day following the recording of their baseline sleep-wake. The baseline was recorded after prior acclimatization of tethering to the recording set-up for at least four days.
*2. Opto-pain mode:* In this model, we used CGRP-ChR2 mice generated by crossing CGRP-creER mice with Channelrhodopsin-2 (ChR2)-floxed mice (Ai27D mice, strain #:012567, RRID: IMSR_JAX:012567, The Jackson Laboratory). Ai27D mice express an improved ChR-2/tdTomato fusion protein following exposure to cre recombinase, so these mice can be used in optogenetic studies for rapid *in vivo* activation of excitable cells with blue light illumination (473 nm). Thus, CGRP-ChR2 mice express ChR2- Tdtomato in all CGRP cells and terminals (validation shown in **Fig. S2**). We used these mice to selectively activate CGRP nociceptors in the hind limb, through the pre-implanted miniature devices, using blue laser light directed to the foot pad (NeuroLux, u-iLED, 6mm). Stimulation of the nociceptors was conducted using the NeuroLux wireless system, which creates a radiofrequency field in the cage for the duration of stimulation, so that the stimulation frequency, pulse duration, and time of stimulation can be set remotely without tethering or handling the mice. Sleep-wake behavioral states were recorded in all mice while subjecting them to nociceptor stimulation every 5 mins for 4s, with or without simultaneous inhibition of the PBel^CGRP^ neurons using red laser light through pre-implanted, bilateral, optical fibers in the PBel (635nm). The red laser stimulus was used to inhibit the PBel^CGRP^ neurons and was always preceded by nociceptor stimulation by 10s that lasted for 20s, overlapping and then extending past the nociceptive stimulus. We found that the stimulation strength of the blue laser light at 8Hz (10ms) activated the nociceptors, however, but wakefulness was produced only in 60% of the trials with a latency of 38.5 ± 11.2s, while 10Hz (10ms) for 4s caused arousal in nearly 100% of the trials with significantly shorter latency, so we continued to use the latter frequency for all the subsequent experiments.

### Effect of genetic deletion of the PBel^CGRP^ neurons on AIP-induced sleep loss

To test the role of PBel^CGRP^ neurons awakening mice in response to pain, we conducted these experiments in the CGRP-creER mice [20], as well as their wildtype litter mates that did not express the cre-recombinase enzyme (WT). To confirm the deletion, we immunolabeled the tissue sections for CGRP using an antibody raised against the CGRP peptide (**Fig. 1c**). CGRP-creER mice that were injected with the adeno-associated viral vector expressing the diphtheria toxin subunit A (AAV-Flex-DTA; 150nl) in a cre-dependent manner had bilateral ablations of the PBel^CGRP^ neurons, while similar injections in the WT control mice did not kill the CGRP neurons in PBel but expressed mcherry (**Fig. 1a-c**). The mCherry fluorescent tag expressed with an injection of the AAV-Flex-DTA vector depicts the surviving, non-cre-expressing neurons in the PBel of both CGRP-creER and WT mice, demonstrating the specificity and location of the genetic deletion. Our group has previously validated this transgenic mouse model (17) and genetic deletions using AAV-Flex-DTA [16,17,21].

**Figure 1.**
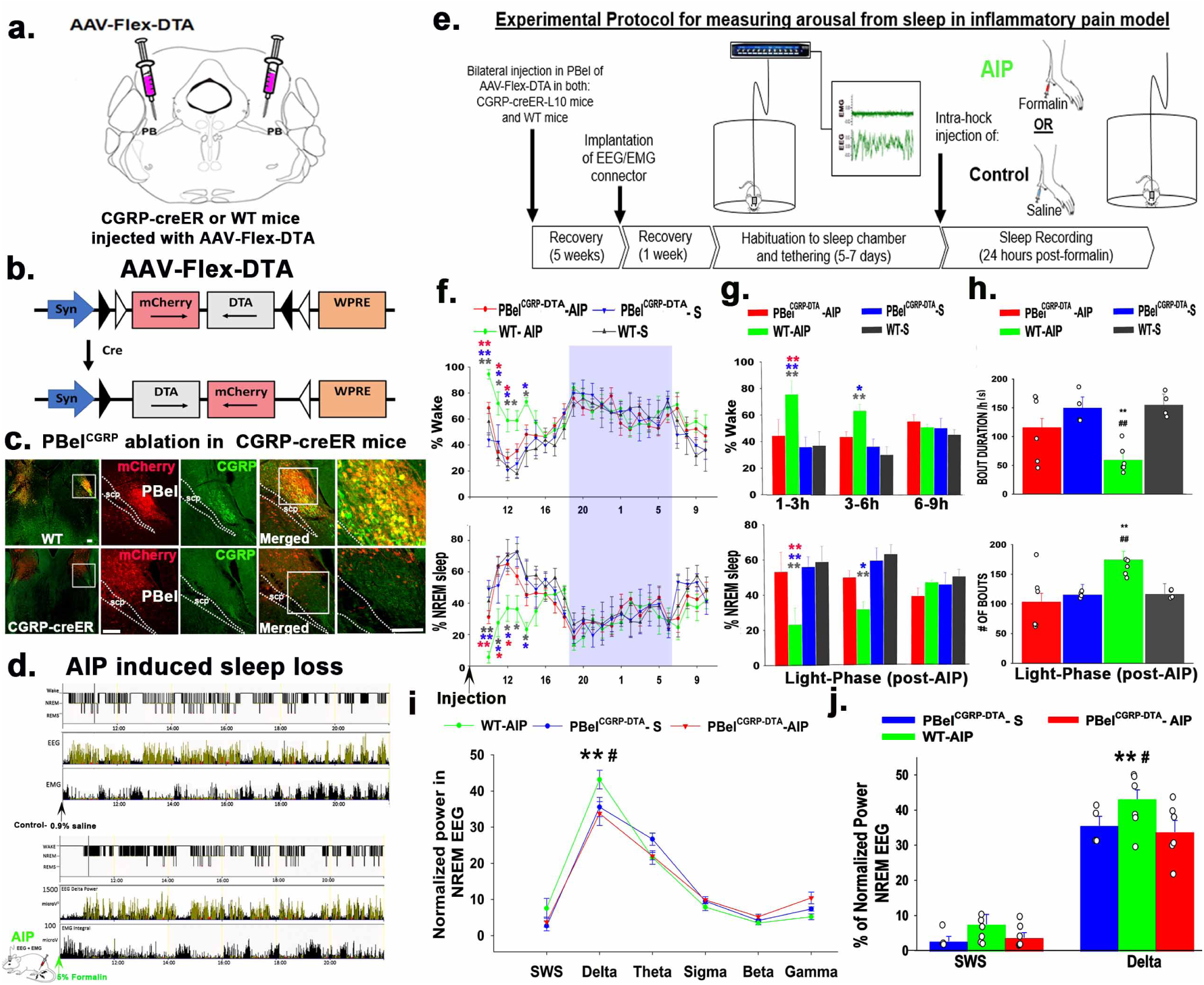
*Genetic deletion of PBel^CGRP^ neurons and sleep in the Acute inflammatory Pain (AIP) model:* CGRP-creER or WT mice were injected bilaterally in the PB with the viral vector AAV-Flex-DTA (**a, b**), which expresses diphtheria toxin subunit A (DTA) in the cre expressing CGRP cells in the PBel. Expression (Cre-dependent) of DTA deletes nearly all CGRP neurons in the CGRP-creER (**bottom panel, c**), but in the WT mice, which lacked cre recombinase, neurons expressed mcherry (**red**) in the intact CGRP (**green**) neurons that were not ablated (**top panel, c**). **Fig. d** shows that representative images of the panels with the hypnograms scored for non-REM (**green**), REM sleep (**red**), and Wake states (**black**) from two treatment groups (**after control**-saline injection or **after AIP**-formalin injection) along with the EEG delta power (**second panel**) and integral EMG (**third panel**). A representative of AIP-induced sleep loss (bottom three panels) is shown by almost no sleep in the first hour and sleep fragmentation in the first 3h, compared to the control above. **Fig. e** represents the timeline of experimental procedures followed for measuring sleep following the AIP protocols. Mean (± SEM) of the percentage of time spent in wake and sleep compared between the treatment groups with the deletions of the PBel^CGRP^ neurons (PBel^CGRP-DTA^) and no deletion (WT) are shown as graphs in (**f**) for every hour for the light and dark phase (blue rectangle). Bar graphs in **g** compare the averages (± SEM) of wake and sleep percentages in the bins of 3h during the light phase post-AIP. Comparison of the sleep bouts numbers/ light phase and average sleep bout durations/h, between the treatment groups are shown as bar graphs (**h**), with data points (representative of n, each mouse) added to each bar. The line graph in (**i**) compares the normalized EEG power during NREM sleep in different frequency bins. Bins of EEG power frequencies during NREM: in SWS (0.5-1.5Hz), delta (0.5-4Hz), theta (5-9Hz), sigma- (10-15Hz), beta (16-29Hz) and Gamma (30-60Hz). Individual data points with bar graphs for SWS and delta frequencies are shown in **J**. The groups were compared using a one-way (treatment) or two-way (treatment X time) ANOVA, followed by Holms-sidak method for multiple comparisons, where **-P<0.001; *-P<0.01. The color of the asterisk in **f-g** represents the comparison group, while in h-j * -represents a comparison to **PBel^CGRP-DTA^-AIP** and #-comparison to **WT-S.** Scale in c: 100μm. Abbreviations: PB- parabrachial subnucleus; PBcl- central lateral PB subnucleus; PBel- external lateral PB subnucleus; scp- superior cerebellar peduncle.

Upon comparing the PBel^CGRP^ neurons in the CGRP-creER deletion and WT control groups, we found that AAV-Flex-DTA induced bilateral deletions in 6 of the 8 injected mice, while the remaining 2 had only unilateral deletions. Both groups were recorded for sleep-wake after injection of saline (control) or 5% formalin in the hind limb (AIP) (**Fig. 1e**). Sleep-wake after saline (S) or formalin (AIP) was then compared between the groups with bilateral PBel^CGRP^ deletions (PBel^CGRP-DTA^; n=6) and those with intact CGRP neurons (WT; n=6). AIP in the WT mice (n=6; WT-AIP) produced significantly higher wakefulness, 69.12 ± 1.2%, compared to 33.64 ± 3.5% in the saline injection PBel^CGRP-DTA^ group (n=6, PBel^CGRP-DTA^ –S), and 35.98 ± 0.28% in the WT saline injection group (n=6, WT-S, F_3, 159_= 13.7; P<0.001) in the first 6 hours post-injection. Correspondingly, the amount of NREM in the first 6 hours post-injection was significantly reduced in WT-AIP compared to WT-S by 53.17% (F_3, 159_= 13.4; P< 0.001) or PBel^CGRP-DTA^-S by 55.12% (P<0.001), before it returned to baseline in the last three hours of the light phase (**Fig.1f and g**). Comparing the number of sleep bouts (period of NREM or REM sleep lasting at least 10 seconds) and amount of NREM during the light-phase (9h) post-AIP suggests that AIP-produced sleep fragmentation causes a significantly higher number of NREM sleep bouts (F_1, 16_= 5.4; P= 0.033) and reduction in average bout duration (F_1, 16_= 14.0; P= 0.002) compared to both saline injection groups (**Fig. 1h**). In addition to sleep loss and fragmentation, we also examined the occurrence of sleep spindles (**Fig. S1b**) and found that AIP significantly reduced sleep spindles (F_2,33_= 5.1; P<0.001) during the first 3 hours post-injection. The impact of AIP-induced sleep loss and fragmentation is further illustrated in **Fig. 1d**, which shows representative hypnograms of EEG with delta power and integral values of EMG in one mouse after injection of either saline or formalin, confirming that the effect of AIP is maximal within the first three hours post-injection.

The PBel^CGRP-DTA^ group showed significantly less AIP-induced sleep disruption than the WT group, demonstrating a sleep loss recovery effect of 84.83% in the first three hours (P<0.001) and 89.5% (P<0.001) in the following three hours post-injection during the light phase (**Fig. 1f and g**). PBel^CGRP^ deletion also significantly reversed sleep fragmentation such that both the bout length and the number of bouts were not significantly different compared to the saline injection condition for both PBel^CGRP-DTA^ (P= 0.56) and WT groups (P= 0.96) (**Fig. 1h**). To investigate whether AIP induced sleep-loss causes changes in EEG power, we compared the power frequencies during NREM sleep-delta (0.5-4Hz), and slow wave sleep, a subset range within delta (0.5- 1.5Hz), and found that AIP in the WT-DTA (CGRP intact) condition caused a significant increase in the NREM EEG power only in the delta frequency range (F_2, 91_= 2.38; P= 0.01; **Fig.1i and j**) over the first 3 hours post-injection. However, the increased power was not significant when averaged over 6 hours post-injection. Since there is clearly AIP-induced sleep loss in the first hour, this suggests that in the subsequent 2 hours, there is significant compensatory sleep rebound, as seen by the increase in delta power, despite the persisting decrease in total NREM and increased sleep fragmentation compared to control conditions. Lastly, despite no significant differences between the NREM EEG power within the spindle frequency range (sigma- 9-15Hz; **Fig. 1i**), the WT-AIP (3 spindles/min) group has significantly reduced sleep spindles (F_2,33_= 5.1; P<0.001) compared to WT saline (P<0.001; 7-8 spindles/ min) and PBel^CGRP-DTA^ saline (P= 0.006) injection groups, which showed reversal effects in the PBel^CGRP-DTA^ (P= 0.018; 5 spindles/min) AIP group (**Fig. S1b**).

### Optogenetic silencing of the PBel^CGRP^ neurons during acute opto-pain stimulus

After validating the CGRP-ChR2 mice for the presence of ChR2 (co-expression of mcherry) in both CGRP neurons and their terminals sites (**Fig. S2**), we conducted experiments using a wireless optogenetics system (NeuroLux Inc., Champaign, IL) to activate nociceptors in the foot pad using blue laser light (473nm) (opto-pain model). In this model, we tested the necessity of the PBel^CGRP^ neurons to relay pain signals that cause arousal by acutely silencing these neurons using far-red laser light through bilateral optical fibers acting on the PBel^CGRP^ neurons that expressed an inhibitory opsin, AAV-Flex-JAWS (PBel^CGRP-JAWS^), as per the paradigm showed in **Fig.2a**.

### In-vitro validation of JAWS optogenetic silencing

Using brain slices, we confirmed that the JAWS-expressing PBel^CGRP^ neurons were inhibited by red (635nm) laser light (**Fig. 2c-iii-vi**). Continuous 20-sec red laser light exposure hyperpolarized JAWS- expressing PBel^CGRP^ neurons (-46.70 ± 8.12 mV; n = 6) and silenced the neuronal firing (**Fig. 2c-iii and v**), whereas similar light stimulation was ineffective towards PBel neurons that did not express JAWS (-0.05 ± 0.45 mV; *n* =4) (**Fig. 2c-iv and vi**).

**Figure 2.**
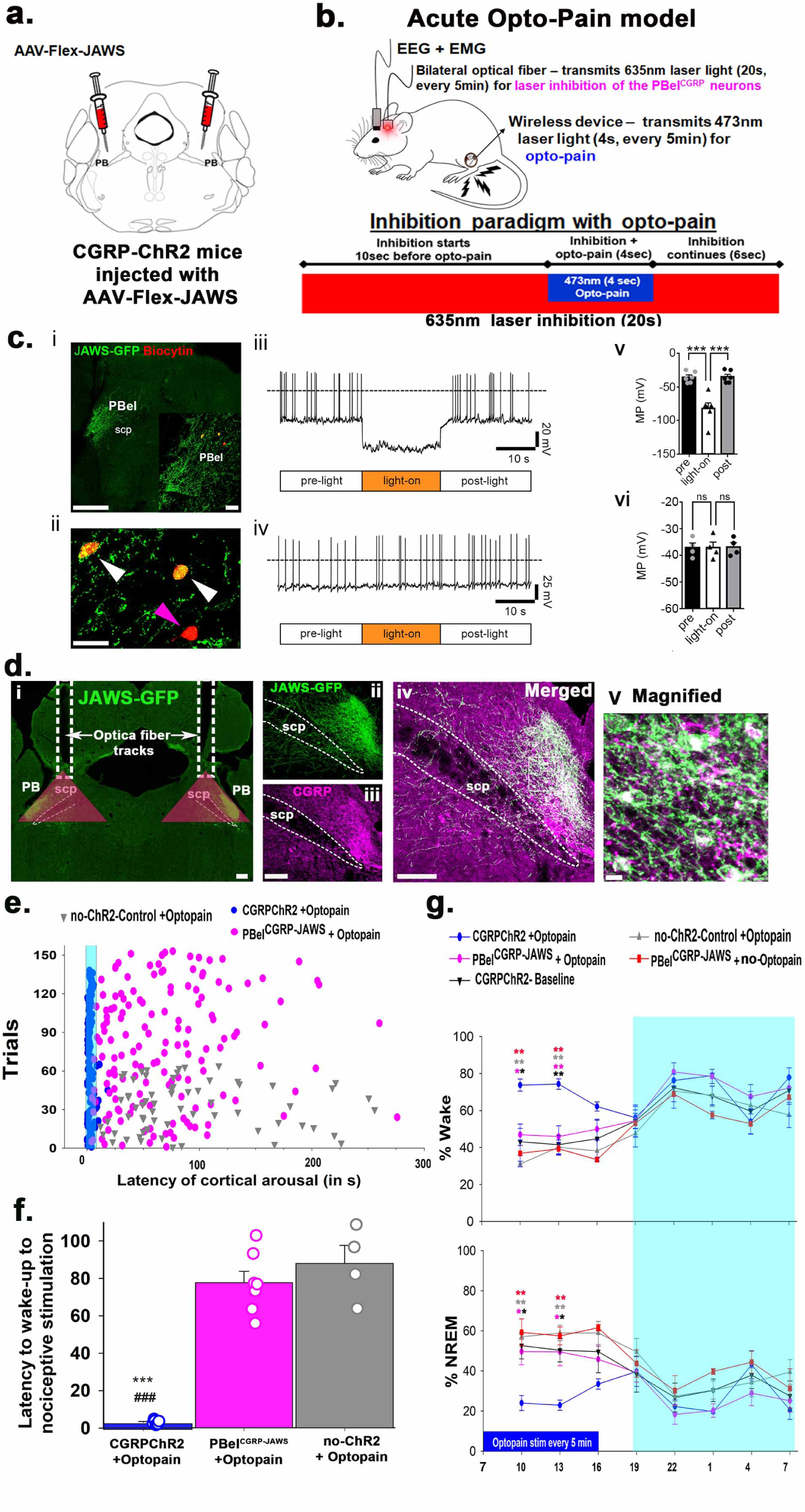
*JAWS mediated photoinhibition of the PBel^CGRP^ neurons in the Opto-Pain model:* In these experiments, we used CGRP-ChR2 mice that expressed ChR2 in all the CGRP neurons and terminals (**validation in SFig1**), which were injected bilaterally with AAV-Flex-JAWS-EGFP in the PB (**a**) and implanted with EEG/EMG, bilateral optical fibers and µLED wireless device in the foot pad for stimulation of the CGRP- nociceptors for the acute opto-pain paradigm (**b**). *Invitro validation of JAWS expressing PBel^CGRP^ neurons* **(c): i**) Examples of recorded brain slices showing JAWS-GFP expression in the PBel (green; scale bar: 500 µm) and *post hoc* labeling of recorded PB neurons filled with biocytin from the recording pipette and then labeled with streptavidin-AF-555 conjugated (orange-red, *insert;* scale bar: 50 µm). **ii**) Image showing the same biocytin labeled PBel neurons at higher magnification (white arrows: two recorded PBel^CGRP^ neurons that express JAWS-GFP; purple arrow: a recorded PBel neuron that does not express JAWS-GFP; scale bar: 50 µm). **iii**) Light exposure (635nm wavelength; 20 s duration) hyperpolarized and silenced action potential firing of PBel^CGRP^ neurons expressing JAWS. **v**) Bar histogram graphs representing the mean membrane potential before (pre), during (light-on), and after (post) light stimulation. One-way ANOVA, *F*: 32.46; *p* < 0.0001, *n* = 6; Bonferroni’s multiple comparisons *post-hoc* test; *** *p* = 0.0001 pre vs light-on; *** *p* = 0.0001, light-on vs post. **iv**) Neurons of the PBel that do not express JAWS do not respond to light. **vi)** The bar histogram graph represents no changes in the mean membrane potential (MP) before (pre), during (light-on), and after (post) light stimulation. One-way ANOVA, *F*: 0.21; *p* = 0.8149, *n* = 4; Bonferroni’s multiple comparisons *post-hoc* test; *p* = > 0.9999 pre vs light-on; *p* = > 0.9999, light-on vs post. PBel, external-lateral parabrachial nucleus; scp, superior cerebellar peduncle. *JAWS expressing CGRP neurons in the PBel* **(d):** Photomicrographs show that AAV-JAWS (green)- as seen by the expression of the enhanced green fluorescent protein (EGFP) were selectively expressed in the CGRP (**purple**) cells in the PBel (**i- iv**). These JAWS expressing CGRP cells in the PBel (**PBel^CGRP-JAWS^**) were targeted by the implanted optical fibers (fiber tracks in **i**) to photo-inhibit by the laser light of 635nm (red laser), and the red triangles mark the area (500^2^µm) maximally affected by the transmitted light that include PBel^CGRP-JAWS^ expressing neurons. The inhibition paradigm by red laser that preceded the activation of the nociceptors (**Opto-Pain**) by 10s is explained in **b**. *Effect of the photo-inhibition of the PBel^CGRP-JAWS^ neurons*: The latency of cortical arousals (**e**) in different trials, in response to the activation of CGRP-nociceptors (Opto- Pain) in CGRP-ChR2 mice (blue) were compared with mice where **PBel^CGRP-JAWS^** neurons were photo-inhibited by red laser light (magenta), and also with those that had no ChR2 expression in CGRP neurons and terminals (no-ChR2 control; grey). The mean (±SEM) of arousal latency in the above groups is shown as bar graphs that are also plotted with average data points from each mouse (**f**). Repeated stimulation (4s every 5 minutes) of CGRP-nociceptors also affected the percentage of time spent in sleep and wake, which were compared between the treatment groups in the graphs shown in **g**. The groups were compared using a one-way (**f**) or two-way (**g**) ANOVA, followed by the Holms-sidak method for multiple comparisons, where **, ###- P<0.001; *- P<0.01. The color of the asterisk in **g** represents the comparison group, while in e * -represents the comparison to **PBel^CGRP-JAWS^+ Opto-Pain** and #- comparison to **no-ChR2 + Opto-pain**. Scale in **d**: 100μm (i-iv); and 15μm in magnified panel. Abbreviations: PB-parabrachial subnucleus; PBcl-central lateral PB subnucleus; PBel-external lateral PB subnucleus; scp-superior cerebellar peduncle.

### In-vivo optogenetic inhibition

CGRP-ChR2 (n= 10) or CGRP-creER (no ChR2- control; n=4) mice were injected with AAV-Flex-JAWS (150nl) bilaterally in the PB.

These mice were then implanted with EEG/EMG, bilateral optical fibers directed to the PB, and miniature probes (NeuroLux, u-iLED, 6mm) in the foot pad to stimulate nociceptors in the foot wirelessly and remotely, avoiding any handling of the mice (**Fig. 2a-b and d**). The optical fiber track placement, as well as the specific expression of JAWS (co-expressing GFP) in the PBel^CGRP^ neurons, were validated and assessed with immunohistochemistry staining as shown in **Fig.2di-v**. Out of the 10 CGRP-ChR2 mice, we found that 7 had more than 80% PBel neurons that were transfected with JAWS and optical fibers placement within 1-1.5mm dorsal to the transfected PBel^CGRP-JAWS^ neurons. Therefore, these 7 cases were included for the sleep and latency analysis data shown in **Fig. 2e-g**. Of the remaining three, one had no transfection of JAWS, and the other two had off-target fiber placement.

We found that stimulation of foot-nociceptors at a frequency of 10Hz (10ms) for 4s duration wakes the mice in 100% of trials with significantly shorter latency of 2.98 ± 0.43s, compared to the control mice without ChR2 (87.91 ± 9.6s; F_7,15_= 73.81; P< 0.001). However, when the nociceptor stimulation was preceded by 10s of silencing the PBel^CGRP-JAWS^ neurons (**Fig. 2b**), arousals were prevented, and the latency of wakefulness was significantly increased (77.63 ± 0.61s). Such silencing in the presence of red light has been validated *in vitro* (**Fig. 2c**). The same optogenetic silencing in the three mice with off-target fiber placement or no JAWS transfection did not prevent arousal to the nociceptive stimulus; it showed a similar arousal latency to those that received Opto-pain without PBel^CGRP-JAWS^ inhibition.

Repeated nociceptive stimulation (10Hz; 4s every 5 minutes) with blue laser (ZT1-ZT10) induced a significant increase in the percent time spent in wakefulness (62.9 ± 12.2%; F_4, 143_= 14.2; P< 0.001) and a decrease in NREM (by 48.2 ± 7.3%; F_4, 143_= 13.78; P<0.001) for the first 6h during the light phase, compared to baseline and no-ChR2 controls. Inhibiting the PBel^CGRP^ neurons with red laser without any nociceptor stimulation (PBel^CGRP-JAWS^ + no opto-pain group) did not change the percent time spent in the wake or NREM, which was comparable to the baseline sleep-wake condition, demonstrating that the PBel^CGRP^ control inhibition is not inducing sleep, but preventing otherwise prompted arousals. However, simultaneous inhibition of PBel^CGRP-JAWS^ neurons using red laser exposure with nociceptor stimulation (PBel^CGRP-JAWS^+ opto-pain) recovered the opto-pain induced NREM sleep loss by 95 ± 1.7% (P= 0.004 in 1-3h, and P=0.002 in 3-6h post-stimulation onset, **Fig. 2g)** and significantly reduced the opto-pain induced wakefulness (P= 0.002 1-3h, and P<0.001 in 3-6h post-stimulation onset).

### Opto-inhibition of the PBel^CGRP^ terminal sites during AIP-induced sleep loss

Next, we tested which of the forebrain terminal fields of PBel^CGRP^ neurons relayed nociceptive information to cause arousal by selectively opto-inhibiting each of the four main terminal fields, the SI-BF, the CeA, the BNST, or the LH. We used a cre-dependent inhibitory opsin virus (AAV-hSyn1-SIO-eOPN3-mScarlet, also called AAV-Flex-eOPN3) that suppresses or blocks the excitatory input from the PBel^CGRP^ neurons when the terminals sites are exposed to the laser light, causing selective silencing of the presynaptic terminal fields, as previously been demonstrated[22–24]. AAV-Flex-eOPN3 provides long-lasting, reversible inhibition of synaptic transmission through direct activity on the presynaptic release apparatus and Ca^2+^ channel function and is an effective tool for long-term inhibition without tissue damage as it only requires brief light pulses at low power[22,23].

Injections of AAV-Flex-eOPN3 were made bilaterally in the PB of CGRP-creER mice (n=37) and WT mice (no cre expression; n= 7), creating PBel^CGRP-eOPN3^ mice that express eOPN3 in a cre-dependent manner in PBel^CGRP^ neurons and their terminal fields, while WT mice had no eOPN3 expression (**Fig. 4a-c**). After recovery, all mice were implanted with bilateral optical fibers targeting one of the terminal fields: the SI- BF (n=11), the CeA (n= 8), the BNST (n= 8), or the LH (n= 8). To validate eOPN3 expression in the terminal fields of the CGRP-creER mice, we immunostained the brain sections for dsRed (mScarlet eOPN3 fluorescent tag), and verified the placement of the bilateral optical fiber tracts. Of the 37 mice PBel^CGRP-eOPN3^ mice, 8 mice had optical fibers targeting the SI-BF, 7 mice the CeA, 7 mice the BNST, and 5 mice the LH. These mice also had more than 80% transfection of the CGRP cells in the PBel, which was confirmed with corresponding transfection in the terminal fields (**Fig**. **4a-c**). We compared each of these groups to the WT group following AIP (injection of 5% formalin in the hind limb) and saline injection, with or without the opto-inhibition of the PBel^CGRP^ terminals as per the paradigm mentioned in **Fig. 3d**. We also investigated which of the terminal sites contributed to PBel^CGRP^ mediated regulation of AIP-induced sleep loss (**Fig. 5a**) and sleep fragmentation (**Fig. 5b**).

**Figure 3.**
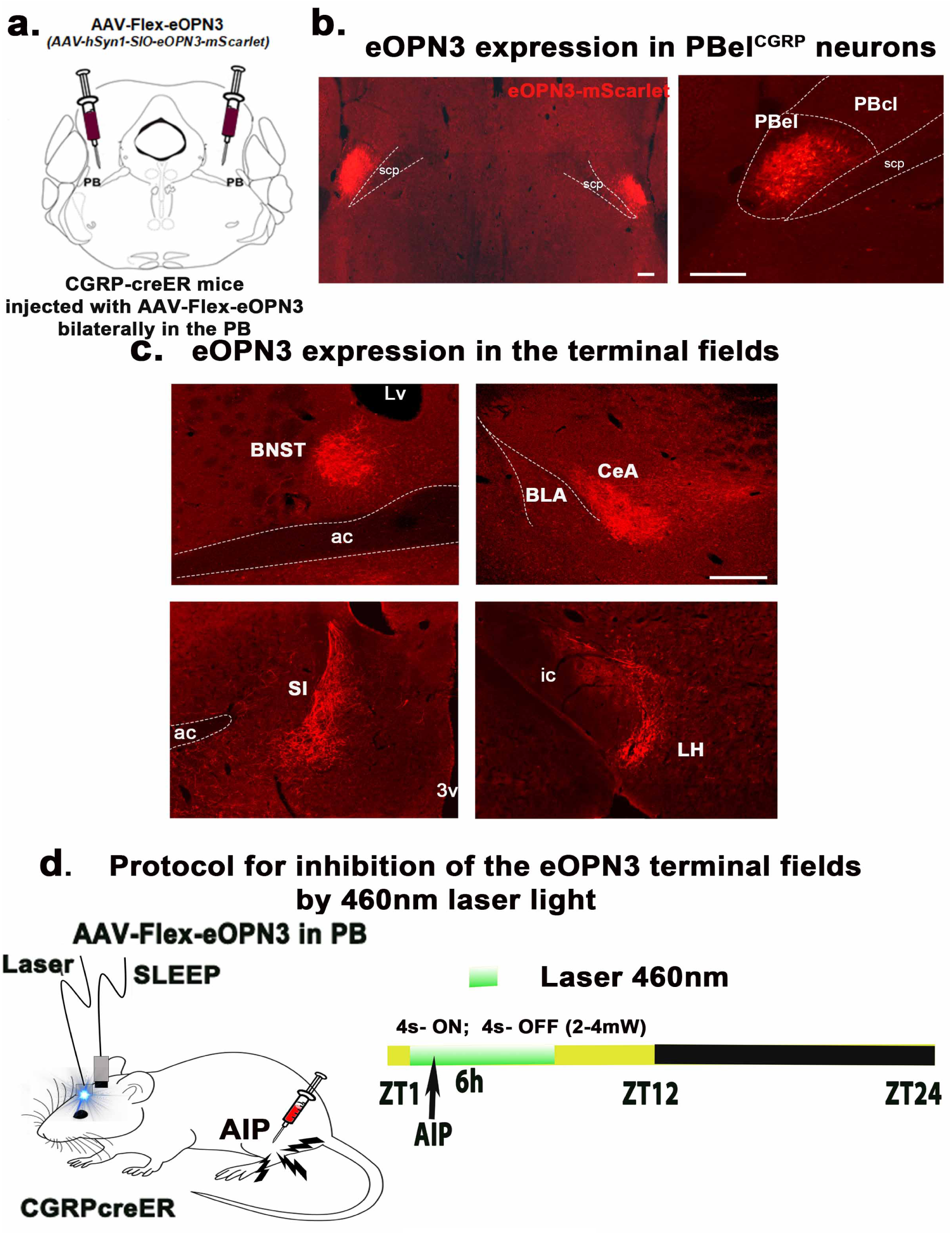
*Expression of eOPN3 in the terminal fields of the PBel^CGRP^ neurons:* CGRP-creER mice were injected bilaterally with the viral vector that co-expresses eOPN3 (AAV-hSyn1-SIO-eOPN3-mScarlet) with mScarlet (immunolabeled with dsRed) in the PBel^CGRP^ neurons (**a**, **b**) and the terminal field (**c**). Scale in b and c- 100μm; Abbreviations: ac- anterior commissure; SI- substantia innominata; BNST- bed nucleus of stria terminalis; ic- internal capsule; CeA- central nucleus of amygdala; LH- lateral hypothalamus; PBcl- central lateral PB subnucleus; PBel- external lateral PB subnucleus; scp- superior cerebellar peduncle. The protocol of optogenetically inhibiting the eOPN3 expressing the terminal fields by the 460nm LED light, which blocks the excitatory input from the PBel^CGRP^ neurons at the targeted site (**d**).

**Figure 4.**
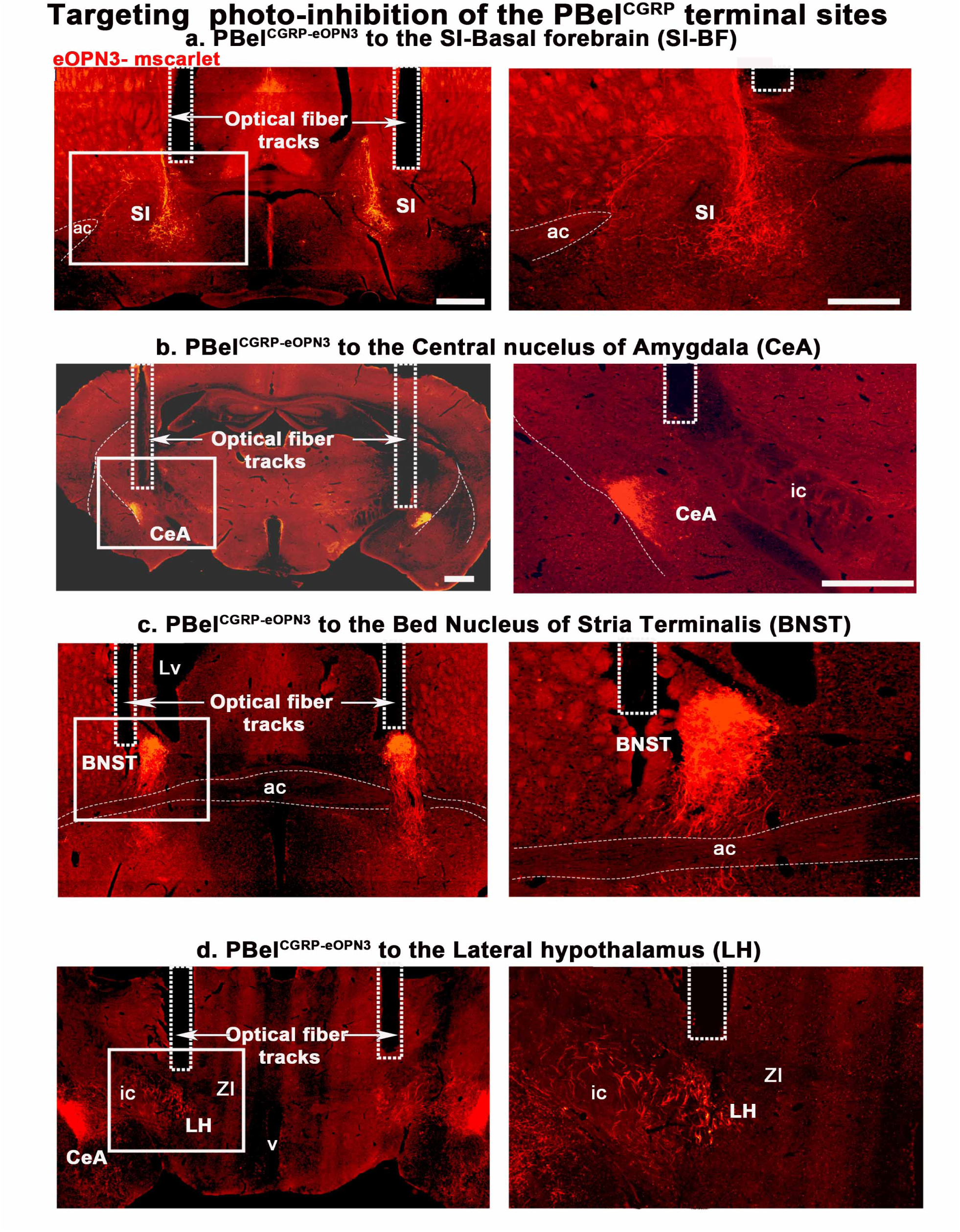
*eOPN3 labeled terminal fields targeted by the optical fibers*: Representative photomicrographs showing the brain sections with the bilateral optical fiber tracks targeting the substantia innominata (SI) in the basal forebrain (BF) (**a**), the central nucleus of amygdala (CeA) (**b**), the bed nucleus of stria terminalis (BNST) (**c**), and in the lateral hypothalamus (**d**) from different groups of mice that were injected with AAV- hSyn1-SIO-eOPN3-mScarlet and immunolabeled for dsRed. Dashed rectangles marked the optical fiber tracks, that illuminated the terminal sites with LED light (460nm) at power of 2-4mW. Scale in a- 500μm and b-d 200μm; Abbreviations: ac- anterior commissure; SI- substantia innominata; BNST- bed nucleus of stria terminalis; ic- internal capsule; CeA- central nucleus of amygdala; LH- lateral hypothalamus.

**Figure 5.**
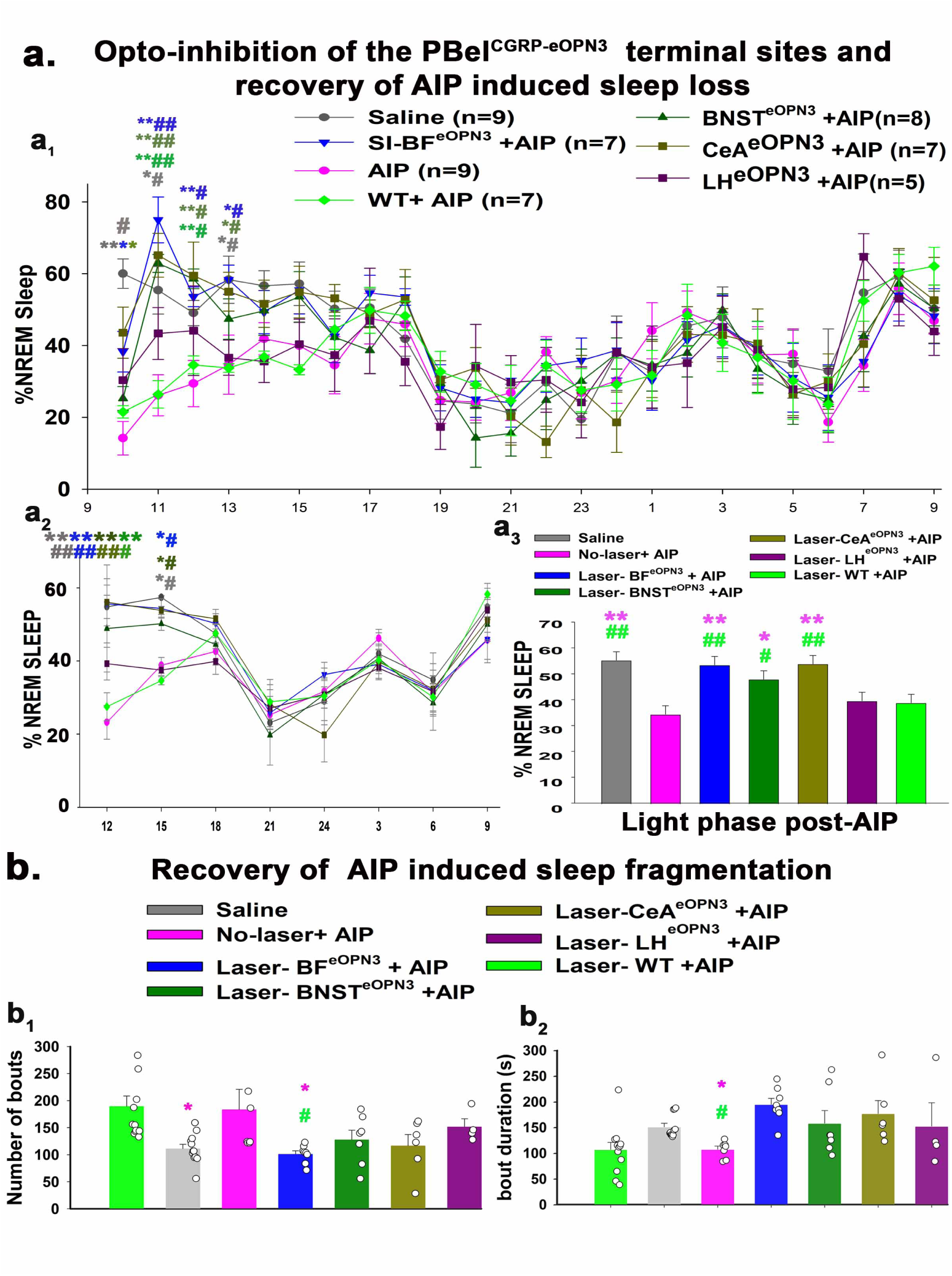
*Comparison of the photo-inhibition of the eOPN3 expressing PBel^CGRP^ terminal fields on recover of AIP induced sleep loss:* The percentage (mean ±SEM) of the time spent in the NREM sleep is compared in the seven treatment groups, where sleep was recorded after either saline (n=9) or formalin injection (AIP; n=9) in the hind limb, with or without photo-inhibition of the terminal fields (**a1-a3**). Of these seven groups, four of them had eOPN3 injected in the PBel^CGRP^ neurons (CGRP-creER mice), and these were implanted with the optical fibers directed to the eOPN3 expressing terminal fields (SI-BF^eOPN3^, CeA^eOPN3^, BNST^eOPN3^, LH^eOPN3^) and recorded for sleep after AIP. Wildtype mice (WT) that had a similar injection of the viral vector in the PB, but it did not transduce the PBel^CGRP^ neurons, and they were also recorded for sleep in response to AIP with laser illumination (WT+AIP; n=7). The graphs in **a1** represent the percentage (mean ±SEM) of the time spent in NREM sleep post-AIP every hour. The data were averaged over 3h bins and shown in **a2**, while the bar graph in **a3** shows the averaged NREM over the light phase post-AIP. Comparison of sleep showed that the order of effect of photoinhibition on sleep reversal was SI-BF> CeA> BNST> LH. The bar graphs with data points representing each mouse in **b** show the comparison of recovery in AIP-induced sleep fragmentation, as seen by the number of NREM bouts (**b1**) and average sleep bout durations (**b2**) in different groups, during the light phase post-AIP. The groups were compared using a two-way (**a1, a2**) or one-way (**a3-b2**) ANOVA, followed by Holms-sidak method for multiple comparisons, where **, ##- P<0.001; *, #- P<0.01. The color of the asterisk represents the comparison of the color group with **AIP**, while # -represents a comparison to **WT+AIP (a1, a2)**. The color of * and # in **a3- b2** represents the comparison to AIP and WT-AIP, respectively.

AIP in WT and PBel^CGRP-eOPN3^ mice (without opto-inhibition) induced significant sleep loss of 45% (F_6,405_= 23.0; P<0.001) in the first 6h compared to saline injection control. AIP also induced sleep fragmentation in the first 9 hours compared to saline injection post-AIP, where there was a significant increase in the number of sleep bouts number (F_6, 49_= 3.9; P=0.003) and a decrease in the bout duration (F_6, 49_= 4.13; P=0.002).

For opto-inhibition of the PBel^CGRP^ terminal sites, blue laser light (460nm) was turned on continuously for 4 seconds, then off for 4 seconds, and then repeated for a duration of 6 hours (ZT2-ZT8) during the light phase following either AIP or saline injection. Both PBel^CGRP-eOPN3^ and WT mice were recorded for four conditions in randomized order: 1) sleep-wake after saline injection, 2) sleep-wake after AIP, 3) sleep-wake after saline injection + 6h of opto-inhibition, and 4) sleep-wake after AIP + 6h opto-inhibition with one-week recovery in between each recording. Of the four terminal sites, we found that PBel^CGRP-eOPN3^ inhibition of SI-BF (SI-BF^eOPN3^) and CeA (CeA^eOPN3^) had very similar recovery of AIP-induced sleep loss (F_6, 147_= 67.42; P<0.001; SI-BF- 98% and CeA- 97.95%), followed by PBel^CGRP-eOPN3^ inhibition of BNST (BNST^eOPN3^; 88.3%; P=0.022) post 6 hours compared to AIP in both PBel^CGRP-eOPN3-^ no laser and WT groups (**Fig.5a2**). Opto-inhibitions of the LH (LH^eOPN3^) showed a trend towards recovery of AIP-induced sleep loss (14.8%), but it was not statistically significant (P=0.85).

The inhibition of terminals at both the SI-the BF and the CeA caused significant recovery in AIP-induced sleep loss as compared to both WT-AIP and PBel^CGRP-eOPN3^- no-laser-AIP (no opto-inhibition) and WT-AIP throughout the first 4 hours post-injection for (**Fig. 5a1;** F_6,405_= 5.9; P<0.001). Terminal inhibition in the BNST caused significant recovery in AIP-induced sleep loss sleep recovery for the second (P<0.001) and third (P<0.001 and P=0.002, compared to AIP and WT-AIP) hours post-injection. Averaging sleep across 3 hours bins post-injection (**Fig. 5a2**) also showed that both SI-BF and CeA had comparable AIP-induced sleep loss recovery (F_6,112_= 3.55; P= 0.003), which was significant for first (P<0.001) and second (P=0.028- SI-BF; P=0.0035-CeA) 3h bins, while AIP-induced sleep loss recovery in the BNST was only significant for the first 3h bin (P<0.001 and P=0.008). These results suggest that PBel^CGRP^→ SI-BF + CeA is presumably the pathway that causes awakenings to pain, with a smaller contribution of PBel^CGRP^ projections to the BNST, and, finally, PBel^CGRP^ projections to the LH contributing the least amount.

### Pharmacological blocking at PBel^CGRP^ terminal sites during AIP-induced sleep loss

We used local pharmacological blocking to understand if pain-induced arousal stimulus is relayed through the CGRP or the glutamatergic NMDA receptors, as all PBel^CGRP^ neurons are glutamatergic [16] and their terminal sites express both CGRP and NMDA receptors[25–27]. For this, we implanted bilateral guide cannulas (26G, Plastics One) in C57bl/6J mice targeting PBel terminal fields in the SI-BF (n=10) and CeA (n=8). We chose these two sites because opto-inhibition at these PBel^CGRP^ terminal sites was most effective in preventing pain-induced arousals. For the pharmacological blocking, we used CGRP receptor blocker injections, BIBN (BIBN4096BS; 80μg/ml; Boehringer Ingelheim Pharma KG, Biberach, Germany) and NMDA receptor blocker injection, AP-5 (40μg/ml), which were delivered using the injector cannula that was placed in the guide and extending past the tip of the guide by 1mm. The total injection volume was 200nl on each side and was administered simultaneously at 3.3nl/sec using the infusion/withdrawal programmable dual syringe pump (Harvard Apparatus, MA, US). These mice were then recorded for six conditions in randomized order: 1) sleep-wake after saline injection, 2) sleep-wake after AIP, 3) sleep-wake after BIBN, 4) sleep-wake after AP-5, 5) sleep-wake after AIP + BIBN, 6) sleep-wake after AIP + AP-5 with one-week recovery in between each recording. After all recordings were completed, all mice received bilateral injections of thionine (50nl) using the same equipment for histological confirmation of injector cannula placement (**Fig.6b**). Of the 18 total mice, 6 mice implanted with bilateral cannulas in the SI-BF, and 5 mice implanted with bilateral cannulas in the CeA had cannula track and injector tip marks (indicated by thionine stain) within 1-2mm in the vicinity of the corresponding terminal sites (as seen by eOPN3 staining in **Fig. 4a-d**), and these were identified in these mice based on the anatomical landmarks (as shown in **Fig.6b**). The remaining 7 cases, where the cannula tips were 3-4mm away from the terminal sites were considered anatomical controls, where the injections of the blockers did not prevent

### AIP-induced sleep loss

AIP alone induced significant sleep loss of 57.6% (F_7, 128_= 50.75; P<0.001) in the 1-3h period and 32.4% (P=0.047) in the subsequent 3-6h period post-injection (**Fig. 6c**) compared to saline injection. We then compared the sleep-wake of the blocking-AIP conditions, with either the saline injection condition or blocking alone conditions. BIBN injections (CGRP antagonist) recovered AIP-induced sleep loss in the SI-BF implanted mice by 97.4%, and 93.8% in the CeA implanted mice in the 3-6h period post-injection. We found that the effects of the NMDA receptor blocker (AP-5) were comparable to the BIBN at both injection sites during the 1-3h and 3-6h periods post-injection. Injections of either BIBN or AP-5 alone (no AIP) in the SI-BF mice did not result in an increase of NREM, such that the sleep-wake amounts post-injection were comparable to the saline condition, demonstrating that the receptor blockers do not specifically promote sleep (**Fig. 6c**), but instead blocked the relay of pain stimulus that leads to wakefulness. Our results also suggest that PBel^CGRP^ neurons may act on both the CGRP and NMDA receptors at the SI-BF and CeA sites, where CGRP may potentiate the effect of glutamate by acting on NMDA receptors [25].

**Figure 6.**
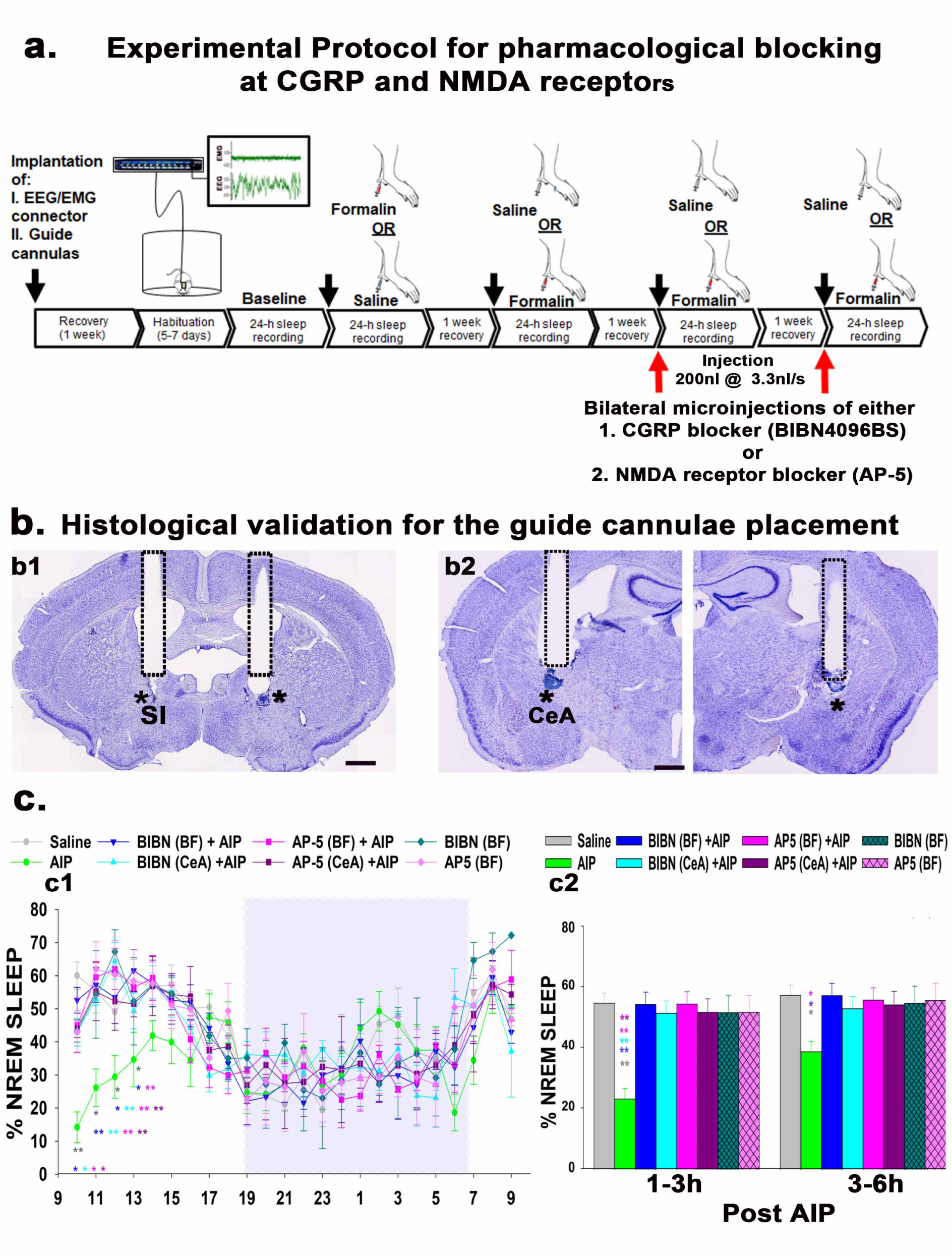
*Pharmacological blocking of CGRP or NMDA receptors in the SI-BF or CeA during AIP-induced sleep loss:* The experimental protocol for recording sleep with or without AIP and after blocking of either the CGRP (BIBN4096BS; 80µg/ µl) or NMDA (AP-5; 40µg/ µl) receptors with 200nl of antagonists injected bilaterally in either the SI- BF or the CeA using the injectors cannulas and the infusion/withdrawal programmable dual syringe pump (**a**). Photomicrographs of Nissl’s-stained brain sections (**b**) show the guide-cannula tracks used to insert the injector cannulas, whose tips were marked using thionine stain, in the vicinity (within a radius of 1-1.5mm) of the substantia innominata (SI, **b1**) in the basal forebrain (BF) and the central nucleus of the amygdala (CeA, **b2**). Asterisks mark the tip of the injector cannula that extended the guide cannulas by 1 mm, observed either in the same or adjacent section that lie within 1mm of the PBel^CGRP^ terminal sites. Scale in **b1 and b2** is 0.5mm. Graphs in **c** show the percentage (mean ± SEM) of time spent in NREM sleep for each hour (**c1)** and in 3h bins (**c2**) after AIP. The blue rectangle represents the dark phase. The groups were statistically compared using a two-way (treatment X time) ANOVA, followed by Holms-sidak method for multiple comparisons, where **- P<0.001; *- P<0.01. The color of the asterisk compares AIP to the group representing the color in the graph.

## Discussion

In the present study, we tested the hypothesis that PBel^CGRP^ neurons act as a relay node of nociceptive stimulus that mediates pain-induced wakefulness. Our findings, using two acute pain models that both reliably induced objectively quantifiable sleep loss, show a reduction in sleep spindle density and increased sleep fragmentation that lasted for 3-6h, providing significant insights into the mechanisms of pain-induced sleep disruption.

In the field of pain research, the formalin administration test (AIP) in mice or rats is a valid and reliable model of tonic pain and nociception and is often used for assessing the analgesic potential of various compounds [28–33]. A subdermal injection of formalin in the hindlimb produces a typical biphasic time course of pain behaviors lasting at least an hour. The first phase of resulting pain behavior is due to direct stimulation of nociceptors in the hindlimb. The second and longer-lasting phase is mediated by a combination of peripheral input and spinal cord sensitization that depends on the extent of peripheral inflammation. This response is unlike that with the TRPV1 receptor agonist capsaicin, which does not activate the dorsal horn and has only shorter and more mild nociceptive responses[34,35]. Therefore, the more prolonged tonic pain induced by formalin is more consistent with acute pain from trauma and injury and is more likely to induce sleep changes seen in patients with acute peripheral injuries [36–40].

Some studies have investigated how sleep fragmentation increases pain sensitivity[6,7,41–43], and others have linked inflammation with sleep disruption[44–49]. Still, none have attempted to objectively investigate if acute pain in AIP models reliably produces quantifiable sleep loss and fragmentation, which can be used to detect analgesic efficacies of various compounds and identify pathways that modify this pain-induced sleep loss. Therefore, we first characterized sleep-wake responses in the AIP model during the early light phase by objectively recording and analyzing sleep-wake and comparing them with control conditions. Our results suggest that AIP resulted in a robust and consistent reduction of sleep by 45-50% over the first 6h, with the most significant reduction occurring in the first hour (by 89%), which also matches the timing of heightened pain behaviors observed in rodents[34,35]. This period was followed by a significant increase in NREM delta power (0.5-4.0 Hz) in the 2-3h post- AIP to recover a near total sleep loss from the 1h and is suggestive of a sleep-rebound. We also observed significantly fragmented sleep that lasted for the remainder of the light-phase post-AIP (9h), which is consistent with observations in other pain models of neuropathic injury[5]. Lastly, despite no change of the EEG power in the spindle frequency range (sigma-10-15Hz), we found that AIP also significantly reduced sleep spindle density for 1-3h post-AIP, which aligns with other studies in rodents[50] and humans[51,52] that suggest spindle density negatively correlates with pain intensity and sensitivity.

Additionally, we used an opto-pain model that involved precise and selective activation of peripheral nociceptors and primary afferents that express CGRP in the hindlimb via laser light (10Hz, 10ms) that produced brief arousals with short latency compared to the control condition, suggesting that the arousals were selective to the activation of the CGRP-expressing primary afferents. Repeated stimulations of nociceptors in the opto-pain model simulated acute peripheral pain-induced awakenings, which reduced sleep (by 48.2%) for a 6h stimulation period and was consistent with the AIP model. Many other labs have produced similar pain models[53–58], but only a few objectively analyzed them for sleep and sleep fragmentation[5,7].

Reliable and quantifiable pain assessment is crucial to its diagnosis and effective management, which can be done with specific biomarkers. Our data from both pain models shows that we could reliably use objective criteria of sleep loss and fragmentation as a tool to detect how manipulations of the PBel^CGRP^ neurons and their terminals affect sleep-wake, therefore comparing efficacies of blocking pain to recover the sleep loss.

Our results of genetically ablating PBel^CGRP^ neurons show that such deletions reversed AIP-induced sleep loss by 87%, reversed sleep fragmentation, prevented the increase in slow wave perception, and increased spindle density, suggesting that there was a significant decrease in nociceptive signals that drive circuits regulating wakefulness [51,52]. Similarly, optogenetic silencing of PBel^CGRP^ neurons (using JAWS) before activation of the CGRP nociceptors to induce pain-like effects also prevented short latency awakenings and completely reversed the opto-pain-induced sleep loss. Results from both the AIP and opto-pain models suggest that PBel^CGRP^ neurons are the key relay through which acute peripheral injury drives wakefulness from NREM sleep.

Our data also corroborates the anatomical findings that the PB receives spinal nociceptive-specific projections from lamina I[10] and is activated by a wide range of nociceptive stimuli [8–11,59]. Activation of excitatory neurons in the lateral PB of naïve mice caused place aversion, while their inhibition reduced pain-related responses [13]. In particular, the PBel^CGRP^ neurons are involved in several components of pain behavior, including escaping from a noxious stimulus and forming aversive memory.

Studies have shown that mice with ablation of PBel^CGRP^ neurons have attenuated reactions to a noxious heat source and do not develop conditioned place aversion, while their activation promotes aversion[60]. Additionally, both neuropathic and inflammatory-induced pain produced amplified responses in these neurons that last in parallel with the increased pain metric measurements (19) for 5 weeks. More recently, ablation of the PBel^CGRP^ neurons has also been shown to attenuate sleep fragmentation in a chronic neuropathic pain model[5]. Similarly, in our study, mice with either genetic deletion or optogenetic silencing of PBel^CGRP^ neurons had almost no sleep loss in response to pain stimulus, suggests that blocking this node attenuates pain-induced disruption of normal sleep. However, this is not due to their sleep-promoting effect, as inhibition alone without a pain stimulus did not increase or promote sleep. Our data also reinforces results from our previous study, which showed that these neurons regulate cortical arousals in response to other aversive stimuli, like hypercapnia (brief periods of pauses in breathing- apnea^17,55^ during sleep), through their projections to CeA and SI-BF.

Next, we dissected the underlying neural pathways by conducting optogenetic inhibition of four different terminal sites to understand which are critical for driving cortical arousals from sleep in response to primary afferents in the AIP model.

Optogenetic blocking of the terminal sites followed the effects order of SI-BF > CeA > BNST > LH for reversal of AIP-induced sleep loss and fragmentation. As shown by others[8,15,19], our results also confirm the importance of the PBel^CGRP^ → CeA pathway in pain-induced sleep disturbance and, in addition, identified the novel roles of two other projection sites, SI-BF and BNST, which have long been associated with mediation of arousal[62–66] and stress-related fear and anxiety[27,67–69].

Additionally, our data is supported by a recent study, which shows that cholinergic neurons in the nucleus basalis area of the BF that receive projections from the PB are overactive in a mouse model of neuropathic pain [70]. Similarly, there are other studies suggesting that BNST mediates nocifensive responses to pain[15]. However, in contrast to some of the previous studies[69,71], we observed that the PBel^CGRP^→ LH pathway did not have a significant role in mediating pain-induced sleep disturbances, which could be attributed to sparse projection patterns. These results parallel with our earlier study, where we investigated the arousal response to hypercapnia[16] and found that inhibition of the PBel^CGRP^ terminal field in the BF had the most profound effects, while inputs to the LH made little or no contribution.

Lastly, pharmacological blocking at either the SI-BF or CeA using CGRP or NMDA receptor antagonists had proportionally similar effects on reversing AIP-induced sleep loss. PBel^CGRP^ neurons are glutamatergic, so their activation in response to pain causes the release of both glutamate and CGRP, which then act on NMDA and CGRP receptors at their terminal sites. Furthermore, our findings align with an earlier study that investigated the LPB→CeA circuits in slice recordings and showed that synaptic transmission is mediated by glutamate [25,72]. However, the fibers from the LPB also release CGRP in the CeA, where it acts to potentiate synaptic NMDA-R function[25,69], which could have a potent impact on the strengthening of the nociception-emotion-sleep link in persistent pain. Both NMDA or CGRP blockers injected in SI-BF selectively blocked the acute-pain-induced sleep disturbances but did not alter normal spontaneous sleep, analogous to an earlier study where neither cell nor vesicular glutamate transporter (Vglut2) gene deletions in the LPB changed spontaneous sleep [61]. A recent study also identified the role of PBel^CGRP^ in chronic neuropathic pain-induced microarousals [5]. Thus, our results help identify the precise role of the PBel^CGRP^ pathways that cause awakenings in response to nociceptor activation during acute tonic pain, which may be clinically relevant in preventing pain-induced sleep disturbances without affecting normal spontaneous sleep.

In conclusion, our findings establish a neural network that links pain and sleep disturbances. This process is mediated by the transmission of nociceptive signals through the spino-parabrachial pathways, which converge on the PBel^CGRP^ node in the brainstem[10]. The glutamatergic PBel^CGRP^ neurons play a role in regulating these sleep disturbances by projecting to key forebrain arousal areas, namely the substantia innominata of the basal forebrain (SI-BF), and the central amygdala (CeA), in a similar proportion. These neurons also project to the bed nucleus of the stria terminalis (BNST). At these target sites, their effects are mediated by action on NMDA and CGRP receptors. These results suggest that these sites could be targeted for the development of non-addictive analgesics, utilizing compounds that act on non-opioid receptors found in these areas. Recently, CGRP receptor blockers (known as gepants) and monoclonal antibodies targeting CGRP receptors have effectively treated headaches and migraines [73–75], but the central sites of action for these treatments are unclear. It is unlikely that they cross the blood-brain barrier and target CGRP receptors in the BF or CeA at concentrations that allow them to alter neuronal responses [76–78]. However, whether high central CGRP-receptor antagonism may provide any additional therapeutic benefit or help the non-responders [79] remains to be understood.

The bidirectional link between sleep and pain potentially involves the same anatomical pathways, where prolonged sleep disturbances may alter pain sensitivity, decrease the effectiveness of opioid analgesics, and delay recovery from pain [6,7,41–43]. Therefore, assessing sleep disturbances in both acute and chronic pain could serve as a useful biomarker for evaluating the effectiveness of therapeutic interventions aimed at managing pain and enhancing patients’ quality of life.

Future research should adopt an integrative approach that combines objective assessments of sleep and pain. This can involve utilizing existing or novel interventions designed to specifically target and investigate common anatomical pathways that can improve the reduced quality of life (i.e., disease burden) due to sleep deprivation in chronic and acute pain patients. Additionally, future studies could leverage innovative tools, such as a spatially resolved transcriptional atlas[80], to identify specific cell types and unique receptor types located at key points along these pathways that could be targeted to alleviate both sleep disturbances and pain.

## Methods

### Animals

Male mice (CGRP-creER, CGRP-ChR2, their wildtype littermates, and C57BL/6J mice) were bred in our Animal Research Facility using a heterozygote breeding scheme to ensure true littermate as control animals. All mice were backcrossed to the C57BL6 strain for at least six generations and were outbred to C57BL/6J mice obtained from Jackson Labs every three generations. Mice were group-housed until implantation, then singly housed with ad libitum access to water and food, ambient temperature of 21–23°C, humidity levels between 40 and 60%, and a 12-hour light/dark cycle. Wildtype littermates were randomly assigned to experimental groups. All animal procedures and protocols were approved by the Beth Israel Deaconess Medical Center Institutional Animal Care and Use Committee and met National Institutes of Health standards.

### Validation of mice

CGRP-ChR2 were validated by double immunolabeling for CGRP and mcherry (co-expressing ChR2) (**SFig. 2**), where the neurons in PB that were CGRP were labeled for ChR2, and similar labeling was also seen in terminal sites, e.g., in the CeA and BNST, suggesting reliable expression of ChR2 in the CGRP neurons and terminals. CGRP-creER mice have been validated by us for the eutopic presence of the enzyme cre-recombinase in the CGRP-expressing neurons [16].

### Viral vector

We previously validated AAV-Flex-DTA for specific cre-dependent ablation of PBel^CGRP^ neurons in the CGRP-creER and DR-serotonergic neurons in the Sert-Cre mice [16,17,21]. AAV8-hsyn-Flex-JAWS-GFP (obtained from University of North Carolina Vector Core, Lot # AV6640B, 3.2x10^12^ virus molecules/ml dialyzed w/350mM NaCl & 5% D-sorbitol in PBS) has also been verified by our group and others for causing effective inhibition by red laser (635nm) / LED light **(Fig.2c**). Additionally, we used pAAV5-hsyn1-SIO-eOPN3-mScarlet-WPRE (Addgene, Catalog # 125713-AAV5, Lot # v128152), which has also been validated by others for causing effective inhibition with blue (460nm) light [22–24].

### Surgery

*For genetic deletion, optogenetic silencing and opto-inhibition experiments:* Under surgical anesthesia, male CGRP-creER or CGRP-ChR2 transgenic mice aged 6-12 weeks and their wild-type littermate (as controls) received micro-brain injections (320nl) of a viral-vector (AAV-Flex-DTA, AAV8-hsyn-Flex-JAWS-GFP, or pAAV-hSyn1-SIO- eOPN3-mScarlet) targeted bilaterally to the PB area (AP: -5.2 to -5.3mm; DV- 2.7 to - 2.8mm; ML: ±1.3 to 1.33mm). To induce cre-expression, mice received intraperitoneal injections of tamoxifen (75mg/kg diluted in corn oil) for five consecutive days concurrent to their corresponding micro-brain injections. After 4 weeks of recovery, all injected mice were implanted with EEG and EMG electrodes. Mice that received AAV-Flex-JAWS were also implanted with bilateral optical fibers (Doric Lenses, dual fiber optic cannulas, 200µm core, 2.6mm distance between fibers, 3 to 5mm length, TFC_200/245-0.37_5mm_TS2.6_FLT) targeted to the PB (same coordinates as above), and a NeuroLux micro-LED (u-iLED) inserted sub-dermally with the flexible micro-fiber directed to the footpad (hind paw) and secured under the skin with sutures (NeuroLux implantable devices, 473nm, 4mm). In addition to EEG/EMG electrodes, mice that received AAV-Flex-OPN3 were also implanted with bilateral optical fibers targeted to one of the terminals sites, SI-BF (AP: -0.15mm; DV -4.5mm; ML: ±1.2mm), BNST (AP: +0.3mm; DV -4.0mm; ML: ±1.0mm), CeA (AP: -1.35mm; DV -4.4 to -4.6mm; ML: ±2.75mm), or LH (AP: -1.4mm; DV -5.0mm; ML: ±1.4 to 1.5mm). Mice were singly housed following implantation and allowed 7-10 days of recovery.

#### For pharmacological blocking experiments

Under surgical anesthesia, male wild-type C57BL/6J mice aged 6-12 weeks were implanted with EEG and EMG electrodes as well as cannulas targeted to either SI-BF (AP: -0.15mm; DV -4.5mm; ML: ±1.2mm) or CeA (AP: -1.35mm; DV -4.4 to -4.6mm; ML: ±2.75mm). Mice were singly housed following implantation and allowed 7-10 days of recovery.

### Data acquisition

Following surgical implantation, mice were habituated to the recording chambers, and requisite tethering equipment used to record EEG and EMG for 7-10 days. Additionally, mice recorded for optogenetic silencing and opto-inhibition were allowed at least 2 days to also habituate to the fiber optic cables. All recordings were acquired at least 5 weeks post-injection of viral vectors to ensure optimal expression and started at ZT2.

Each mouse was connected to a preamplifier, which then connected to a commutator, and then an 8242: 3-channel analog adapter, all from Pinnacle Technology. The analog adapters were fed into a data acquisition console from Cambridge Electronic Design (Cambridge, United Kingdom) and recorded in their corresponding software, Spike2.

### Genetic deletion of the PBel^CGRP^ neurons during AIP induced sleep loss

Recording all the treatment groups for 24 hours of the EEG and EMG recordings after the following conditions were conducted in a random order with 7 days recovery between each: 1) no intervention (baseline), 2) following 25μl 0.9% saline injection in the hind limb or 3) following 25μl 5% formalin sub-dermally in the hind limb (AIP).

### In vitro electrophysiological recordings of the PBel^CGRP^ neurons that expressed JAWS for validating silencing by Laser

For *in vitro* electrophysiological recordings, we used CGRP-creER mice (n=4). We injected 200nl of AAV-FLEX-JAWS-GFP into the lateral parabrachial nucleus (PB) of the CGRPCreER mice, using the coordinates described above. Three to four weeks after AAV injections, the mice were deeply anesthetized with isoflurane (5% in oxygen) via inhalation and trans-cardially perfused with ice-cold ACSF (N-methyl-D-glucamine, NMDG-based solution, as described below). The brains were then quickly removed and coronally sectioned at 250 µm thickness in ice-cold NMDG-based ACSF using a vibrating microtome (VT1200S, Leica). We first incubated the slices containing the PB in the NMDG-based ACSF for 10 minutes at 37°C, then transferred them into a holding chamber at 37°C containing Na-based ACSF for 10 minutes. The brain slices were allowed to gradually return to room temperature (∼1 hour) before recording. Brain slices were submerged in a recording chamber and perfused with Na-based ACSF (1-1.5 ml/min, described below). We recorded PBel^CGRP^ neurons that expressed JAWS-GFP (identified by green fluorescence) and those in the PB that did not express JAWS-GFP as controls (no fluorescence), using a combination of fluorescence and infrared differential interference contrast microscopy (IRDIC).

Recordings were made using a fixed-stage upright microscope (BX51WI, Olympus America) equipped with a Nomarski water immersion lens (Olympus 40X / 0.8 NAW) and an IR-sensitive CCD camera (ORCA-ER, Hamamatsu), with real-time imaging acquired using Micro-Manager software. Neurons were recorded in current-clamp configurations using a Multiclamp 700B amplifier (Molecular Devices), a Digidata 1322A interface, and Clampex 9.0 software (Molecular Devices). Neurons showing more than a 10% change in input resistance over the course of the recording were excluded from the analysis. We photo-inhibited the JAWS-GFP expressing neurons in PB using full-field light openings (∼ 1.8 mW/mm2, 1mm beam width) from a 445 mW LUXEON light-emitting diode (#M595L3; Thorlabs, Newton, NJ, USA) coupled to the epifluorescence pathway of the microscope. We recorded the PB neurons before, during, and after 20s-long continuous light activation. We recorded using a K-gluconate-based pipette solution, in whole-cell current clamp mode (described below). In all the recordings, we added 0.5% biocytin to the pipette solution to mark the recorded neurons [81]. After *in vitro* recordings, to label the recorded neurons filled with biocytin, we fixed the recorded slices in 10% buffered formalin, washed them, and incubated them overnight in streptavidin-conjugated Alexa Fluor 555 (1:500; Cat#: S32351; Invitrogen, Thermo Fisher Scientific Waltham, MA). We acquired images using a Leica Stellaris 5 confocal microscope using a 63X oil immersion objective [82].

### Solutions for electrophysiological recordings

NMDG-based ACSF solution containing (in mM): 100 NMDG, 2.5 KCl, 1.24 NaH_2_PO_4_, 30 NaHCO_3_, 25 glucose, 20 HEPES, 2 thiourea, 5 Na-Lascorbate, 3 Na-pyruvate, 0.5 CaCl_2_, 10 MgSO_4_ (pH 7.3: 95% O_2_ and 5% CO_2_; 310-320 mOsm). Na-based ACSF solution contained (in mM): 120 NaCl, 2.5 KCl, 1.3 MgCl_2_, 10 glucose, 26 NaHCO_3_, 1.24 NaH_2_PO_4_, 4 CaCl_2_, 2 thiourea, 1 Na-L-ascorbate, 3 Na-pyruvate (pH 7.3-7.4 in 95% O_2_ and 5% CO_2_; 310-320 mOsm). K-gluconate-based pipette solution containing (in mM): 120 K-Gluconate, 10 KCl, 3 MgCl2, 10 HEPES, 2.5 K-ATP, 0.5 Na-GTP (pH 7.2 adjusted with KOH; 280 mm). We purchased all other chemicals from Fisher Scientific (Waltham, MA) or Sigma-Aldrich (Saint Luis, MO).

### Optogenetic silencing of the PBel^CGRP^ neurons during acute opto-pain stimulus

All treatment groups were recorded for the 24 hours of EEG and EMG recordings under the following conditions, which were conducted in random order with 7 days of recovery between each treatment: 1) no intervention (baseline), 2) Opto-pain (10hz, 10ms, 4 seconds on every 5 minutes from ZT1-ZT10, 473nm) without inhibition of PBel^CGRP^ neurons, 3) Opto-pain + opto-inhibition stimulation (20 seconds-on and 5 minutes-off from ZT1-ZT10, using red laser- 635nm), where laser inhibition preceded the opto-pain stimulation by 10 seconds for each trial.

Opto-pain stimulation was triggered with the NeuroLux Optogenetic Smart System via the PDC box and auto-tuner with external loop antennae, which were placed around the recording chambers (NeuroLux, Northfield, IL, USA). This allowed for activation of the radio frequency field generated by the antennae to trigger the implanted NeuroLux device, causing the emission of 473nm laser light within the device implanted in the hindlimb (for which the µLED was directed to the footpad) that optogenetically stimulated CGRP peripheral receptors, which caused local pain sensation. The optogenetic silencing of PBel^CGRP-JAWS^ neurons was triggered via a stimulation protocol executed by Spike2 that was also used to simultaneously record EEG and EMG for sleep-wake. The red laser light was transmitted through a fiber optic splitter for bilateral stimulation (TM105FS1B, Thorlabs, NJ) to fiber optics cables (1.5m length, 200µm diameter core, Doric Lenses, Quebec, QC, Canada). The laser power emitted from the end of the fiber optic cables was measured before and after recordings to ensure it was within a range of 8-10mW, though this is likely a higher estimate due to light lost at the connection of the fiber optic cable and implant. We have previously used laser stimulations at similar power for comparable and longer durations (continuous stimulation for 1 minute every 5 minutes) (17), yet we have not observed any tissue damage, as is also the case for the present experiments.

### Opto-inhibition of the PBel^CGRP^ terminal sites during AIP- induced sleep loss

All groups of mice undergo 24 hours of EEG and EMG recordings, under the following conditions that were conducted in random order with 7 days of recovery between each: 1) no intervention (baseline), 2) following 25μl 0.9% saline injection in the hind limb, 3) following 25μl 5% formalin (AIP) in the hind limb 4) following saline with opto-inhibition (4 seconds on, 4 seconds off from ZT2-ZT8, using LED of 460nm wavelength), 5) following AIP with opto-inhibition (4 seconds on, 4 seconds off from ZT2-ZT8). LED light stimulation was triggered-ON via script in Spike2, which is used for simultaneous sleep- recordings. LED (460nm) light was transmitted (Prizmatix, Holon, Israel) through a fiber optic coupler (Fiber Optic Couplers, 1×2 FOC /200u/L3.0M/1.25/FC, Splitting Ratio: 50:50. 0.37 N.A., Prizmatix) that split the beam into two for bilateral stimulation (4 seconds on, 4 seconds off from ZT2-ZT8, using LED of 460nm wavelength). LED power was measured before and after recordings to ensure it was within the 4-6mW range, though this is likely a higher estimate considering some light is lost at the connection of the fiber optic cable and implant.

### Pharmacological blocking of the terminal sites

All mice implanted with EEG/EMG and guide cannulas targeting either SI-BF or CeA were recorded for 24 hours of EEG and EMG after the following conditions that were conducted in varying order, all with 7 days of recovery in between each: 1) no intervention (baseline), 2) following 25μl 0.9% saline injection in the hind limb, 3) following 25μl 5% formalin (AIP) in the hind limb, 4) following saline injection after bilateral injections of CGRP-receptor blocker (80µg/µl dissolved in 1% DMSO) in either SI-BF or CeA, 5) following AIP after CGRP-receptor blocking, 6) following saline injection after bilateral injections of NMDA-receptor blocker (40µg/µl dissolved in 0.9% saline) in either SI-BF or CeA, 7) following AIP after NMDA-receptor blocking, 8) following CGRP-receptor blocking, and 9) following NMDA-receptor blocking. Receptor blockers were delivered using the injector cannula (33G, Pinnacle Technologies, Roanoke, VA) directed via pre-implanted guide cannulas (C235GS-5-2.0/SPC Guide 39622 26GA DBL 5mm PED, P1 Technologies, Roanoke, VA) at a rate of 100nl/minute for a total volume of 200nl using PE10 tubing (Fisher Scientific) connected to a Hamilton syringe (Catalog # 80330, 701RN 10µl SYR 26s/2”/2, Lot # 1055245).

### Data analysis

#### Sleep analysis

EEG and EMG recordings were converted from Spike2 to SleepSign software (Kissei Comtec, Matsumoto, Nagano, Japan) and analyzed in 4-second epochs for sleep state (wake, non-REM, and REM). The files were first auto-scored using an algorithm based on EMG power, EEG delta, and EEG theta power, then manually scored by researchers blind to the treatment groups (NL, RLS, SS, JDL, SK). EEG power spectral densities (PSDs) were computed using the multi-taper method (Chronux toolbox; http://chronux.org) as described previously[83]. Sleep spindles were detected using an automated algorithm (MATLAB) validated and published earlier[84,85]. Briefly, EEG data was band-pass filtered (10–15 Hz, Butterworth Filter), and the root-mean-squared (RMS) power was calculated to provide an upper envelope of the data. The RMS data was then exponentially transformed to further accentuate spindle-generated signals over baseline. Putative spindle peaks (example in **SFig.1a**) were identified in transformed data via crossing of an upper-threshold value, set as 3.5x the mean RMS EEG power across all states for each mouse. Additional detection criteria included a minimum duration of 0.5 s, based on the crossing of a lower threshold set at 1.2x mean RMS power, and a minimum inter-event interval of 0.5 s.

This automated spindle detection algorithm has been rigorously tested and compared with manual spindle detection[84,85].

### Data and statistical analysis for electrophysiological recordings

Recording data was analysed using Clampfit 10 (Molecular Devices) software. Figures were generated using Igor Pro version 6 (WaveMetrics), Prism 7 (GraphPad, La Jolla, CA), and Photoshop (Adobe) software. We calculated membrane potential changes by comparing values before, during, and after light stimulation. We represented data as mean ± SEM, and *n* refers to the number of cells, and compared group means using one-way ANOVA repeated measures followed by Bonferroni’s multiple comparisons *post-hoc* test. Values showing *p* < 0.05 were considered significant.

### Histology

After all *in vivo* recordings were completed, mice were deeply anesthetized and transcardially perfused initially with 0.9% saline, followed by 10% buffered formalin. Subsequently, brains were collected and stored in 30% sucrose in 0.1M phosphate buffer with 0.02% sodium azide overnight before they were sectioned with a freezing microtome in four series of 30µm sections. Sections of each brain were stored in 0.1M phosphate buffer and 0.02% sodium azide at 4°.

### Immunohistochemistry

For mice recorded in either genetic deletion, optogenetic silencing, or opto-inhibition experiments, one series of sections from each brain was immunolabeled for the proteins using the primary and secondary antibodies described in Table 1. For the genetic deletion experiments, brain sections from the CGRP-creER or WT mice that received brain injections of AAV-Flex-DTA were immunolabeled for dsRed and CGRP (**Fig.1d**) to analyze and confirm PBel^CGRP^ neuronal deletions. For the optogenetic silencing experiments, CGRP-ChR2 and WT mice that received brain injections of AAV8-hsyn-Flex-JAWS-GFP were immuno-stained for GFP and CGRP (**Fig.2d**) to confirm PBel^CGRP-JAWS^ transfection and optical fiber tract location. Lastly, CGRP-creER and WT littermates that were used in opto-inhibition recordings that received brain injections of AAV-hSyn1-SIO-eOPN3-mScarlet were immuno-labeled for dsRed (**Fig. 4**) to confirm if the optical fibers at the terminal sites targeted the transfected terminal fields.

**Table 1:** Details of antibodies used for Immunohistochemistry.

| **<u>Antigen</u>** | **<u>Host</u>** | **<u>Dilution</u>** | **<u>Source</u>** | **<u>Immunogen</u>** | **<u>Specificity</u>** | **<u>Secondary Incubation</u>** | **<u>Dilution</u>** |
| --- | --- | --- | --- | --- | --- | --- | --- |
| CGRP | Goat | 1:1000 | Abcam<br>Catalog #<br>Ab36001<br>Lot #<br>GR253001-9 | - | No staining with genetic deletion of CGRP neurons | Biotinylated anti-goat | 1:200 |
| GFP | Chicken | 1:1000 | Invitrogen<br>Catalog #<br>A10262<br>Lot #<br>2738236 | H3-GFP construct transfected in HEK-293E cells | Absence of staining confirmed in wild type and un-injected mice | Biotinylated anti-chicken alexa fluor 488 (Invitrogen) | 1:200 |
| DsRed | Rabbit | 1:1000 | Takara Bio<br>Catalog #<br>632496<br>Lot #<br>1612022 | DsRed-Express, a variant of Discosoma sp. red fluorescent protein | Absence of staining confirmed in wild type and un-injected mice | Biotinylated anti-rabbit alexa fluora 555 (Invitrogen) | 1:200 |

We followed standard immunostaining protocols that had previously been used by our group [16,17]. All sections were incubated overnight in the respective primary antibody (concentration in Table 1) diluted in phosphate-buffered saline with triton-X (0.25%) and sodium-azide (0.02%). On the second day, sections were transferred to the secondary antibody for 2 hours (concentration in Table 1), which is either biotinylated or had a fluorescent tag (Alex-488 or 555 or Cy5). Those incubated in biotinylated secondary were then incubated with a streptavidin labeled fluorescent (Alexa 555 or Cy5). Finally, sections were mounted on Superfrost slides (Fisher Scientific, Pittsburgh, PA), coverslipped with Dako fluorescent mounting medium (Agilent Technologies, Santa Clara, CA), and then scanned using either Olympus Slideview VS200 slide scanner or Leica Stellaris 5 confocal microscope.

For the mice recorded in the pharmacological blocking experiments, one series of sections from each brain were mounted on Superfrost slides (Fisher Scientific, Pittsburgh, PA), dried overnight at RT, and then stained for Nissl using the following protocol. Slides were first washed in DH_2_0, stained for 1 minute, then in 0.1% thionin diluted in DH_2_0, then dehydrated in varying concentrations of EtOH, cleared in Xylene, coverslipped using Permaslip (Alban Scientific, St. Louis, MO) and then scanned using the Olympus Slideview VS200 slide scanner.

## Acknowledgments

We thank Quan Ha, Rayna Jacob, and Aarohi Gupta for their excellent technical support and help with scoring data.

## Author contributions

NL- experimental design, data collection, analysis, and manuscript writing, RDL, ER, and EA- *In vitro* data collection and analysis, RLS- data collection and analysis, RCT, SS, NG, and JDL- data analysis and brain tissue processing, SSB- mouse breeding program; RB- manuscript preparation and critical insights, ST^-^ EEG power and spindle analysis, manuscript writing, SK- experimental design and conceptualization, data collection, analysis, and manuscript writing.

## Data availability

All data generated to support this study’s findings can be made available as a source data file and requested from the corresponding author.

---

**Supplementary Figure 1 (Fig. S1).**
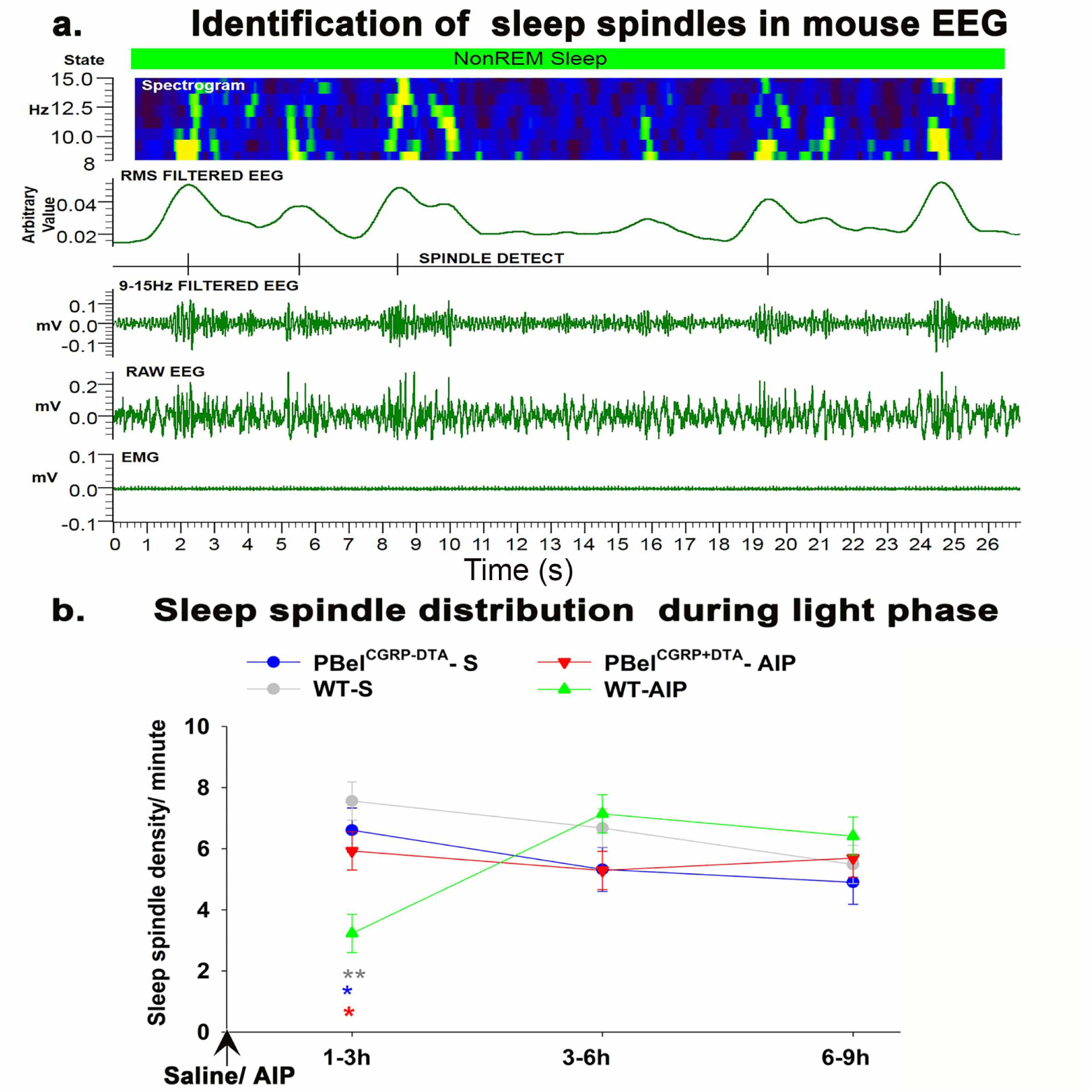
*Depiction of sleep spindles in a representative EEG recording and distribution of sleep spindles across light phase.* A representative photomicrograph of the EEG and EMG recording from a mouse showing the filtered EEG and spectrogram used to identify sleep spindles (**a**). Graph showing the sleep-spindle density (spindles/min) in 3h bins after saline injection or AIP in WT and PBel^CGRP-DTA^ mice (**b**). The groups were compared using a two-way (treatment X time) ANOVA, followed by Holms-sidak method for multiple comparisons, where **- P<0.001; *- P<0.05. The color of the asterisk represents the comparison group.

**Supplementary Figure 2 (Fig. S2):**
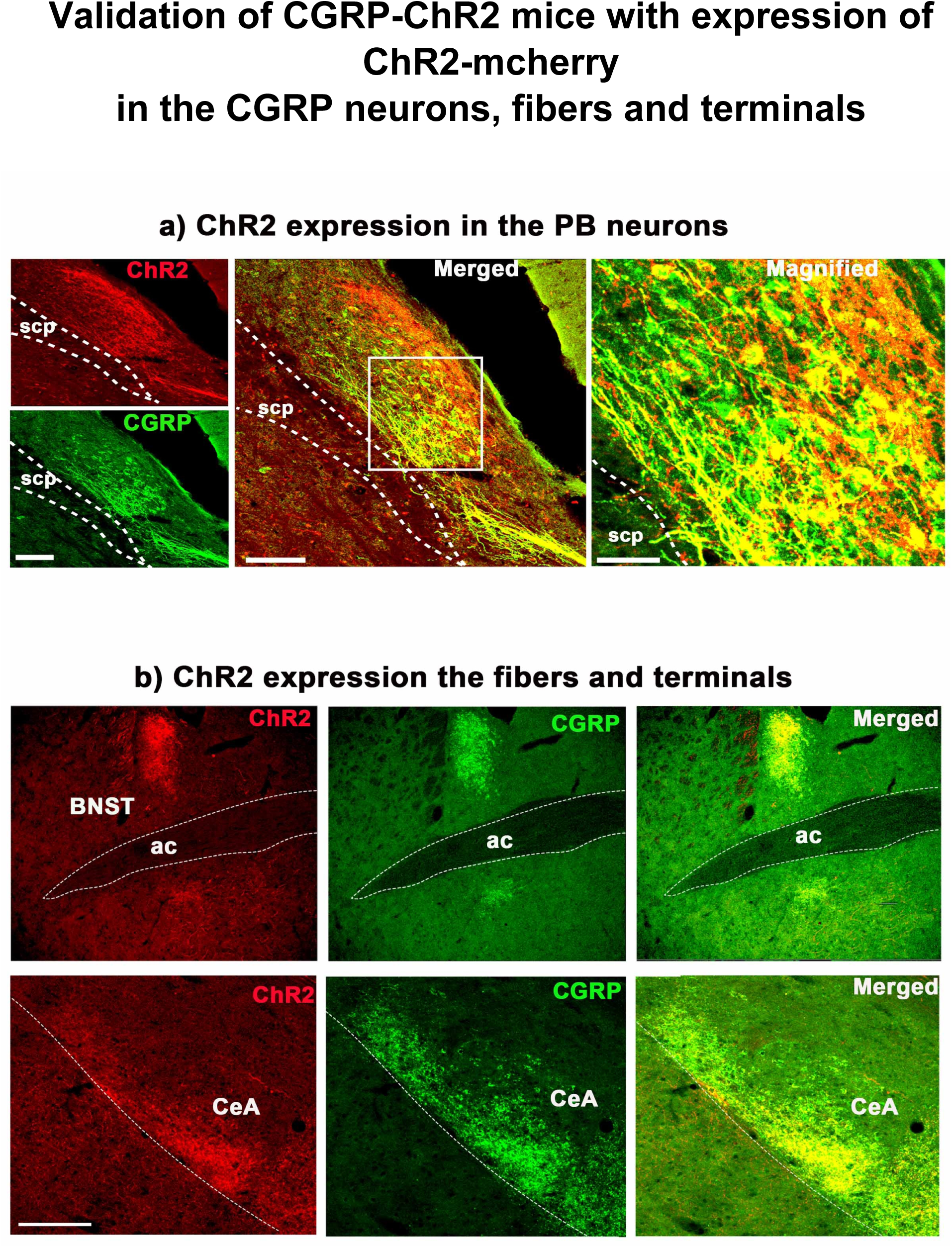
*Validation of Channel Rhodopsin (ChR2) expression in the CGRP expressing neurons and terminal fields in the CGRP-ChR2 mice:* Photo-micrographs showing brain sections from the CGRP-ChR2 mice, immuno- stained for mcherry to label ChR2 (red) and CGRP (green) in the PBel neurons (**a**) and also in the CGRP expressing fibers and terminals seen in BNST and CeA (**b**). Scale: 100µm in a (left) and b; 60 µm and 30 µm in middle and right panels in a.

